# Cooperative recruitment of Yan to paired high affinity ETS sites organizes repression to confer specificity and robustness to cardiac cell fate specification

**DOI:** 10.1101/205724

**Authors:** Jean-François Boisclair Lachance, Jemma L. Webber, Ilaria Rebay

## Abstract

Cis regulatory elements (CREs) are defined by unique combinations of transcription factor binding sites. Emerging evidence suggests that the number, affinity and organization of sites play important roles in regulating enhancer output and ultimately gene expression. Here, we investigate how the cis-regulatory logic of a tissue-specific CRE responsible for *even-skipped* (*eve*) induction during cardiogenesis organizes the competing inputs of two ETS members, the activator Pointed (Pnt) and the repressor Yan. Using a combination of reporter gene assays and CRISPR-Cas9 gene editing, we show that Yan and Pnt have distinct preferences for affinity of sites. Not only does Yan prefer high affinity sites, but a tandem pair of such sites is necessary and sufficient for Yan to tune Eve expression levels in newly specified cardioblasts and to block ectopic Eve induction and cell fate specification in surrounding progenitors. Mechanistically, the cooperative Yan recruitment promoted by this conserved high affinity ETS pair not only biases Yan-Pnt competition at the specific CRE, but also organizes Yan repressive complexes in 3D across the *eve* locus. Taken together our results uncover a novel mechanism by which differential interpretation of CRE syntax by a competing repressor-activator pair can confer both specificity and robustness to developmental transitions.

## Introduction

Development of a multicellular organism relies on tissue-specific gene expression programs to establish distinct cell fates and morphologies. The requisite patterns of gene expression must be both spatiotemporally precise and robust to genetic and environmental variability; this is achieved through the action of transcription factors (TFs), whose activating and repressive inputs are integrated at the cis-regulatory elements or enhancers (CREs) of their target genes. Consequently, the sequence of each CRE provides a physical blueprint for a combinatorial regulatory code that translates upstream signaling information into downstream gene expression. While significant advances have been made in our ability to distinguish regulatory elements from background non-coding genomic DNA and to identify consensus TF binding motifs within them, our understanding of how the intrinsic logic of the cis-regulatory syntax, namely the number, affinity, position, spacing and orientation of binding sites, organizes the necessary set of protein-protein and protein-DNA interactions remains poor (Inukai et al., 2017; Siggers and Gordân, 2014). Because single nucleotide polymorphisms in TF binding sites are being increasingly correlated with altered gene expression and disease susceptibility (Oldridge et al., 2015; Soldner et al., 2016), the ability to deduce the regulatory logic of an enhancer based on its sequence is important.

The tendency for TFs to cluster into superfamilies and for cells to coexpress multiple non-redundant members of the same superfamily implies that enhancer syntax must enable TFs with very similar DNA binding preferences to compete, cooperate and discriminate between binding sites to achieve appropriate gene expression output. Recent insight into these behaviors has come from studies of Hox family TFs (Crocker et al., 2015). The emerging model suggests a specificity-affinity trade-off such that low affinity sites are best discriminated while high affinity sites can be bound by many different Hox factors. Clustering multiple low affinity Hox sites permits the cooperative and additive interactions needed for robust gene activation responses without compromising specificity. Additional work examining Zinc Finger, GATA and ETS family activators in Ciona showed that optimal syntax can compensate for low site affinity to activate strong and specifically patterned gene expression (Farley et al., 2015, 2016). How transcriptional repressors use enhancer syntax to solve the specificity-affinity problem remains to be tested.

The E-Twenty-Six (ETS) superfamily includes both activators and repressors that all recognize the same core DNA sequence, 5’-GGAA/T-3’ (Hollenhorst et al., 2011). ETS TFs are found across metazoan phyla and play key roles in regulating the gene expression programs that direct many aspects of normal development and patterning (Hollenhorst et al., 2011). Exemplifying this, the *Drosophila* transcriptional activator Pointed (Pnt) and the repressor Yan operate downstream of receptor tyrosine kinase (RTK) signaling pathways to orchestrate numerous cell fate transitions (Klämbt, 1993; O’Neill et al., 1994; Rebay and Rubin, 1995; Scholz et al., 1993; Sopko and Perrimon, 2013). Much of the current understanding of Yan and Pnt stems from studying their regulation of *even-skipped* (*eve*) expression during cardiac muscle precursor specification at stage 11 of embryogenesis (Carmena et al., 1998, 2002; Halfon et al., 2000; Knirr and Frasch, 2001) and *prospero (pros)* expression during R7 photoreceptor specification in the developing eye (Hayashi et al., 2008; Xu et al., 2000). Abrogating Pnt-mediated activation or Yan-mediated repression of *eve* or *pros* leads to respective loss or ectopic induction of the associated cell fate (Halfon et al., 2000; Hayashi et al., 2008; Webber et al., 2013a; Xu et al., 2000). Gel shift assays using probes from *eve* or *pros* CREs revealed that most Yan-bound ETS sites are also bound by Pnt (Flores et al., 2000; Halfon et al., 2000; Xu et al., 2000) and subsequent high-throughput assays confirm their preferences for very similar sequences (Nitta et al., 2015; Zhu et al., 2011). Because none of the in vitro biochemistry has been done with full-length proteins, how accurately the results will predict the outcome of Yan-Pointed competition for ETS sites in CREs in vivo is uncertain.

Hints that binding site syntax might influence Yan recruitment come from *in vitro* binding studies with TEL1, the human counterpart of *Drosophila* Yan, and from mathematical modeling of Yan’s ETS site occupancy. TEL1 and Yan, unlike Pnt or its mammalian counterpart ETS1 (Mackereth et al., 2004; Slupsky et al., 1998), self-associate via their Sterile-Alpha Motifs (SAM) and this homotypic interaction is essential for transcriptional repression in both flies and humans (Green et al., 2010; Qiao et al., 2004; Zhang et al., 2010). Using gel shift assays, SAM-SAM interactions were shown to mediate cooperative binding of TEL1 at paired ETS sites regardless of site orientation at spacing of up to 55bp (Green et al., 2010). A recent theoretical analysis of TEL1/Yan occupancy at equilibrium explains how such cooperative SAM-SAM interactions might promote preferential recruitment to tandem ETS binding sites (Hope et al., 2017). Because neither study examined repressive output, the question of whether preferential and cooperative binding of Yan to closely apposed ETS sites might bias Yan-Pnt competition to permit more complex discrimination of CRE syntax than current models assume remains pressing.

To evaluate how CRE syntax organizes Yan-Pnt competition and transcriptional output, we assessed the impact of mutating the eight putative ETS binding sites identified in the *eve* muscle heart enhancer (MHE) that drives *eve* expression in ten segmentally arrayed clusters of pericardial and muscle cells (Halfon et al., 2000). We find that sites with strong affinity best discriminate between Yan and Pnt, with paired sites showing the strongest bias. Thus mutating a tandem of conserved, high affinity ETS sites significantly elevated or expanded reporter expression driven by MHEs from two divergent *Drosophila* species, consistent with compromised repression. Attempts to restore Yan-mediated repression by engineering de-novo pairs of ETS sites into the MHE emphasized the importance of site affinity and spacing, but not of orientation. Using CRISPR/Cas9 gene editing of the endogenous MHE, we showed that mutation of the high-affinity tandem reduced Yan recruitment not only to the MHE, but also to two other CREs across the *eve* locus. Mesodermal Eve expression was elevated, consistent with compromised Yan recruitment resulting in inadequate repression. In this compromised background, two-fold changes in dose of *pnt* and *yan* that would normally be buffered against were now sufficient to induce specification of ectopic Eve+ cells. We conclude that the high affinity ETS site pair within the MHE plays pivotal roles not just in recruiting Yan repressive complexes to the isolated enhancer, but also in longer-range coordination of transcriptional complex organization and function across the locus. More broadly, we propose that differential interpretation of ETS binding site syntax by Yan and Pnt may endow CREs extraordinary flexibility in compartmentalizing and integrating activating and repressive inputs.

## Results

### High affinity Yan binding sites organize transcriptional repression of the MHE

The 312bp muscle-heart enhancer (MHE) from the *even-skipped (eve)* locus provides a tractable system for exploring how cis-regulatory architecture coordinates the activating and repressive inputs that determine expression. Prior work showed that the MHE integrates spatial and temporal cues from multiple signaling pathways and mesodermal determinants to drive expression in a bilaterally symmetric pattern that matches precisely the segmental pattern of ten three-cell clusters of Eve-positive (Eve+) cells within the cardiogenic mesoderm (Halfon et al., 2000) (Fig. 1A-B and Supplemental Fig. S1A). This isolated enhancer also responds appropriately to loss or gain of the TFs that regulate *eve* mesodermal expression, including Yan and Pnt (Halfon et al., 2000; Webber et al., 2013a) (Fig. 1C-E and Supplemental Fig. S1B). Four of the eight GGAA/T core-containing ETS motifs were already known to bind Yan and Pnt *in* vitro (Halfon et al., 2000). In vivo, mutation of any of these four sites reduced reporter expression in the Eve+ clusters while mutating all four sites abolished expression (Halfon et al., 2000). These results suggest that the role of these four ETS sites in Pnt-mediated activation of MHE expression outweighs their contributions to Yan-mediated repression. How the remaining four ETS motifs contribute to MHE expression has not been explored.

**Figure 1.**
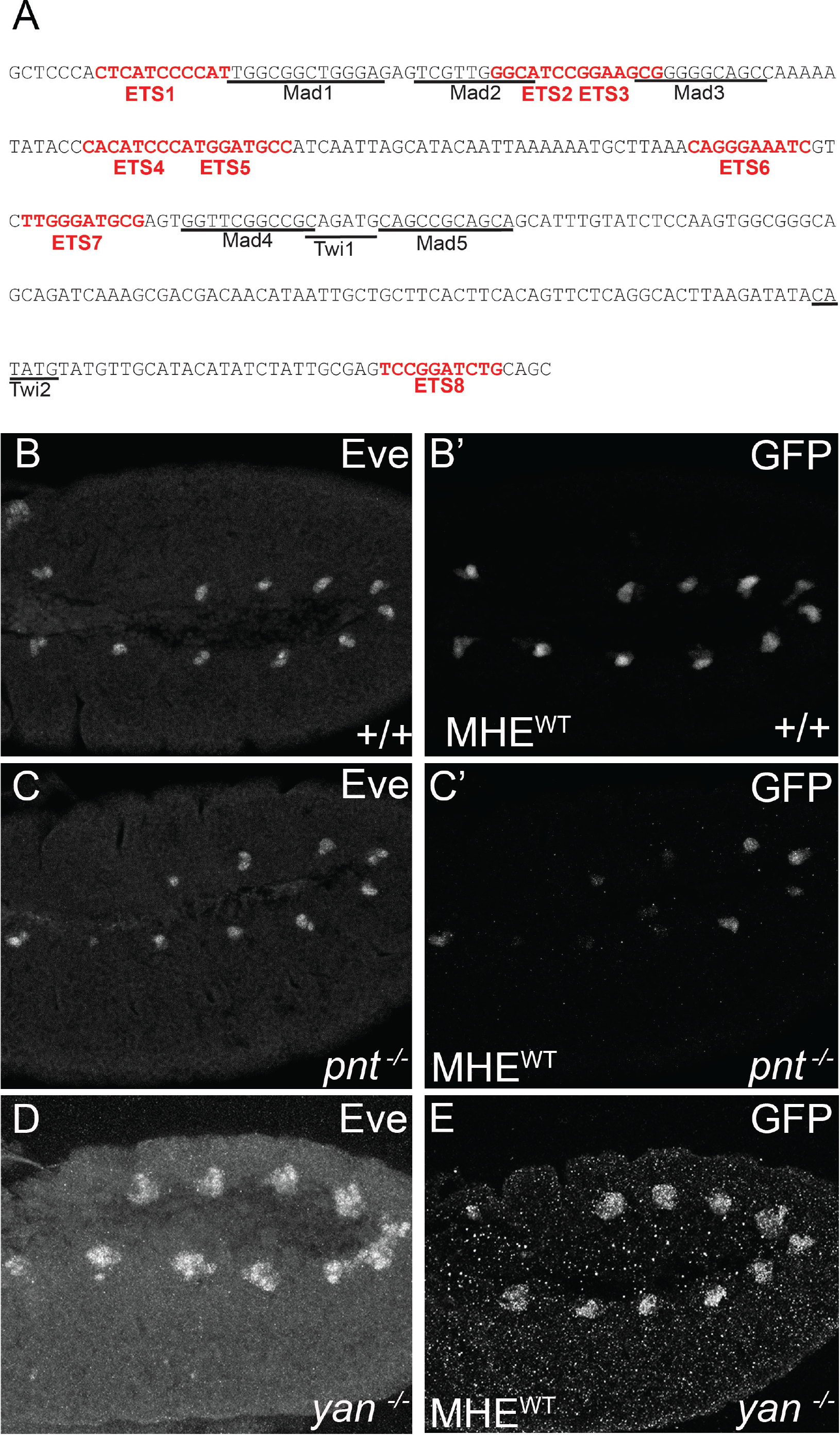
The MHE reliably reports the pattern of mesodermal Eve expression. **A**. Sequence of the MHE with ETS sites in red. Underlining highlights previously characterized Mad and Twi sites (see Supplemental Fig. S1a). **B-D**. Lateral views, oriented anterior to the left and ventral down, of the thoracic and abdominal segments of representative Stage 11 embryos expressing two copies of the MHE^WT^-GFP reporter in *w^1118^* (**B**), *pnt^Δ88^* (**C**) or *yan^ER443^* mutants (**D-E**). Co-staining with anti-Eve (B,C) and anti-GFP (B’, C’) shows that the segmental pattern of cell clusters driven by the reporter matches that of endogenous Eve.

We first compared the affinity of the four putative ETS sites to that of the previously validated sites using a competitive gel shift assay with a labeled consensus ETS sequence and recombinant Yan protein ((Green et al., 2010), see Methods for details). We used Yan rather than Pnt for these experiments both because of the greater sensitivity afforded by Yan’s intrinsically higher in vitro DNA binding ability (Xu et al., 2000) and because Pnt was unstable and prone to degradation. To simplify the nomenclature, we relabeled the sites 1-8 (Fig. 1A). Sites annotated as Ets1-4 in (Halfon et al., 2000) correspond to sites 2, 3, 5 and 8 in our study. Sites 2, 3 and 8 competed as effectively as the consensus ETS sequence, site 6 showed moderate ability to compete, while sites 1,4,5 and 7 exhibited little or no ability to compete (Fig. 2A). All three high affinity sites, but none of the weaker sites, contain C nucleotides at the -1 and -2 positions, in accordance with the binding preferences identified for fly, human and mouse ETS TFs (Nitta et al., 2015; Webber et al., 2013a; Wei et al., 2010). Ranking the eight MHE ETS sites by probability of binding based on Yan or Pnt position weight matrices (PWMs) agrees with the experimental determination of sites 2,3 and 8 as high affinity (Supplemental Tables S1 and S2). The mix of high and low affinity ETS sites made the MHE an ideal test case for assessing how activators and repressors differentially solve the affinity-specificity paradox at target enhancers.

**Figure 2.**
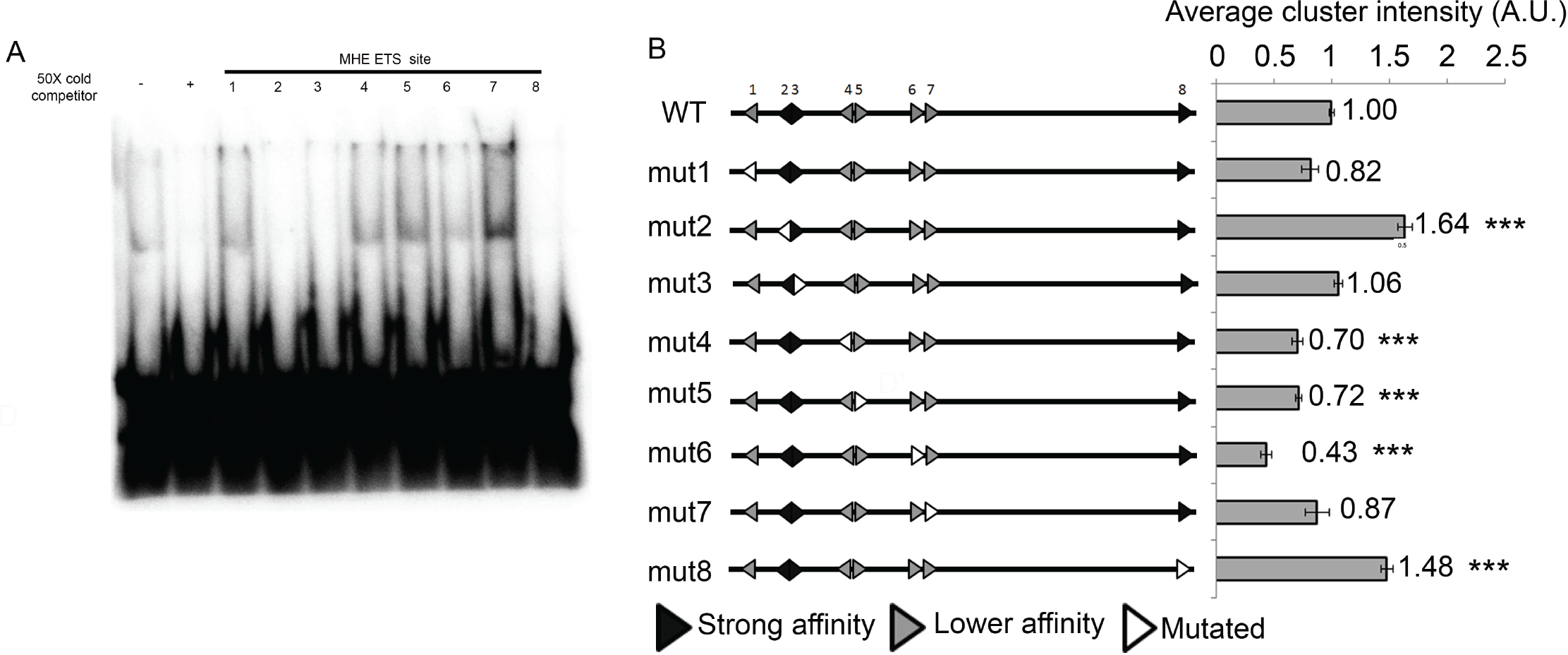
Preferential use of high affinity ETS sites for MHE repression and lower affinity ETS sites for MHE activation. **A**. Competitive gel shift assay shows that MHE ETS sites 2, 3 and 8 compete as effectively for Yan binding as the consensus control, whereas the remaining sites do not. **B**. The impact of individual ETS site mutations on average cluster intensity of the MHE-GFP reporter suggests a role for strong affinity sites in repression and lower affinity sites in activation. In this and all subsequent diagrams of the MHE, ETS sites and their sense/anti-sense orientations are depicted with arrowheads, with lower affinity sites in gray, strong sites in black and mutated sites in white. Error bars show SEM. Statistical significance of single mutants relative to MHE^WT^ after Bonferroni correction is indicated: ***, p <0.001.

To begin, we measured the effects of individual ETS site mutations on enhancer output using MHE-GFP reporter transgenes (Fig. 2B). In broad agreement with Halfon et al (Halfon et al., 2000)., mutation of the majority of ETS sites, namely 1,4,5,6 or 7, decreased reporter expression relative to the wild type MHE-GFP control, with the effect of mutating site 6 the most pronounced. In contrast, mutation of sites 2 and 8 elevated reporter gene expression, while site 3 was neutral. When considered together with the *in vitro* binding data (Fig. 2A), the single-site mutagenesis analysis suggests that high affinity sites (2,3 and 8) contribute more significantly to Yan-mediated repression than lower affinity sites, while lower affinity sites contribute more significantly to activation. Site 3 appeared to be an exception to this rule: it was as effective a competitor as sites 2 and 8 *in vitro*, yet did not alter reporter expression when mutated, leading us to speculate that it might participate in both repression and activation.

To test this possibility, we mutated site 3 together with sites 2 or 8 and compared MHE reporter expression relative to that of the individual mutations. Unexpectedly, simultaneous mutation of sites 2 and 3 induced a greater than four-fold synergistic increase in reporter expression in the Eve+ clusters (Fig. 3A-C) and ectopic expression in surrounding cells (Fig. 3C, white arrows). In addition to defining a role for site 3 in Yan repression, and thereby confirming its functional relevance, the above-additive expression increase within the clusters and the ectopic expression outside the clusters identified sites 2 and 3 as critical cooperative organizers of Yan repressive complexes at the MHE.

**Figure 3.**
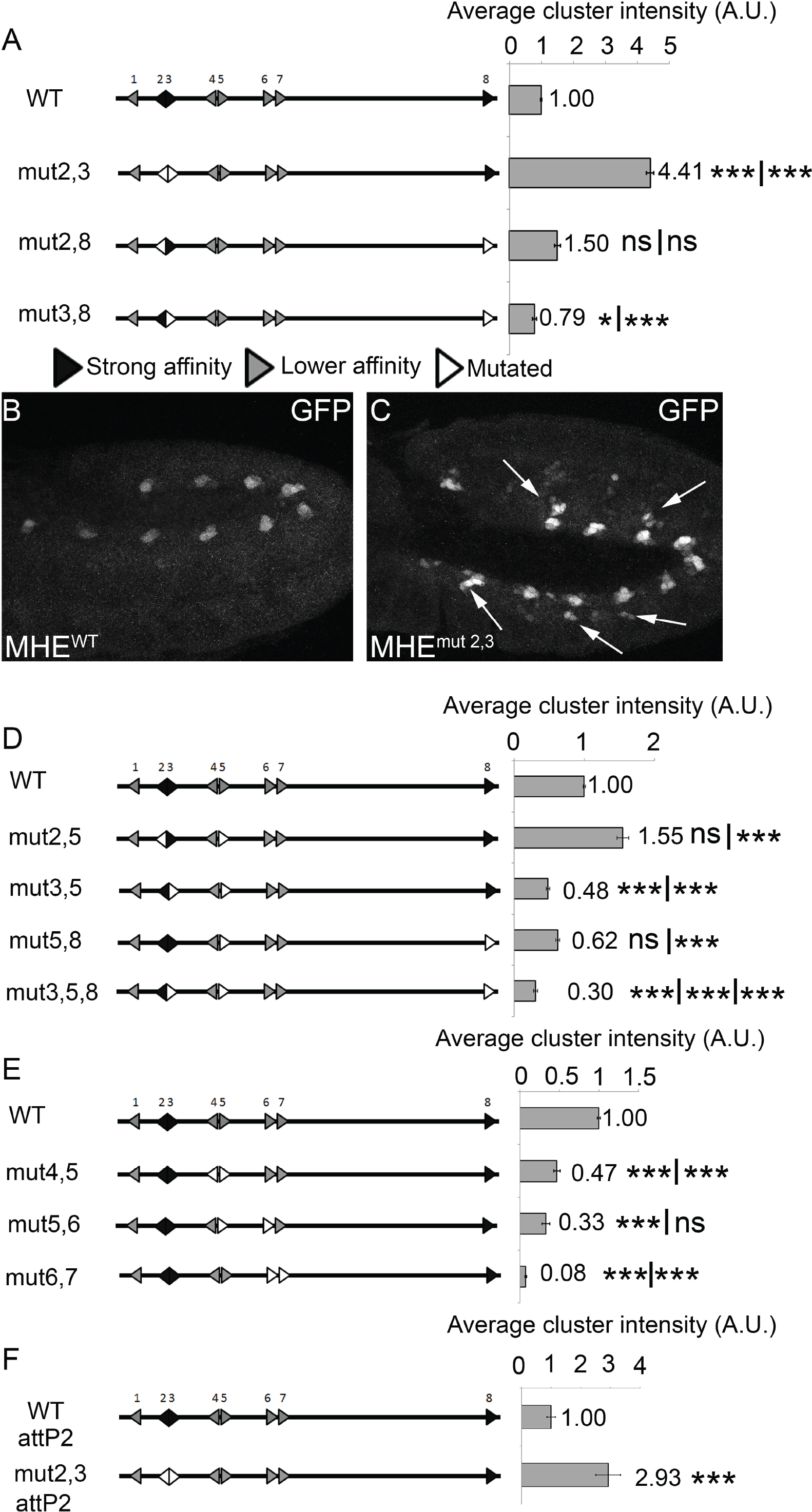
Multi-site mutagenesis reveals key roles for the ETS2,3 pair in limiting expression in Eve+ cells and preventing ectopic activation in surrounding cells. **A,D,E**. Quantification of the impact of mutating pairs of ETS sites on MHE-GFP reporter expression. Statistical significance of double or triple mutants relative to each of the relevant single sites mutations after Bonferroni correction is provided in numerical order: *, p <0.05: **, p <0.01; and ***, p <0.001. **A**. Mutation of the strong affinity 2,3 pair produced a non-additive increase in MHE-GFP reporter expression. Simultaneous mutation of sites 3 and 8 reduced reporter expression indicating that all three high affinity sites also receive activating inputs. **B, C**. Representative Stage 11 embryos showing the increased and ectopic expression driven by one copy of MHE^mut2,3^ relative to MHE^WT^. Arrows highlight ectopic expression. **D**. Mutation of a strong site (2,3 or 8) with the weak site 5 reduced reporter expression, confirming that sites 3 and 8 also contribute to activation. The triple mutant strongly reduced reporter expression, suggesting that MHE activation requires more than five ETS sites. **E**. Mutation of pairs of weaker sites highlighted their primary role in MHE activation, with mutation of the 6, 7 pair causing a greater than additive loss of expression. **F**. Comparison of expression levels driven by MHE^WT^ and MHE^mut2,3^ reporters inserted at a different genomic location (attP2) independently confirmed the results in **A-C** (see Supplementary Figure 1). Error bars show SEM.

In marked contrast, simultaneous mutation of sites 3 and 8 reduced expression below wild type MHE levels (Fig. 3A). This result further confirmed the functional relevance of site 3 and uncovered a role in activating expression. In other words, when site 3 is mutated alone, the lack of expression change reflects balanced repressive and activating inputs; these opposing inputs are revealed only when site 3 is mutated together with sites 2 or 8, respectively. The lower than wild type expression of MHE^mut3,8^ suggests that site 8, despite elevating reporter expression when mutated alone, also mediates Yan-Pnt competition. Consistent with this conclusion, expression of an MHE^mut2,8^ reporter was sub-additive relative to that of the single mutants, although still above wild type levels (Fig. 3A). To summarize, we propose that the high affinity sites 2 and 3 cooperatively recruit and organize Yan-repressive inputs, with Yan-Pnt competition at sites 3 and 8 limiting the strength and stability of repression.

As an independent test of this interpretation, we mutated site 5 together with sites 2, 3 or 8. First, given the results with MHE^mut2,8^, we predicted that loss of a low-affinity site should not change MHE^mut2^ expression; indeed MHE^mut2,5^ showed comparably elevated expression as MHE^mut2^ (Fig. 3D and 2B). Second, just as simultaneous mutation of sites 3 and 8 had reduced MHE expression (Fig. 3A), so mutating a low affinity site in combination with site 3 or 8 should “reveal” the activating roles of those two sites. As predicted, MHE^mut3,5^, MHE^mut5,8^ and MHE^mut3,5,8^ all exhibited significantly reduced expression (Fig. 3D). As controls, we generated three more double mutants to re-confirm the importance of lower-affinity ETS sites to MHE activation (Fig. 3E). In all cases, reporter expression was less than that measured for the single mutants. Simultaneous mutation of site 6 together with weak sites 5 or 7 resulted in the greatest loss of MHE expression among all double site mutant combinations tested.

Finally, we tested how loss of the high affinity ETS sites 2 and 3 impacted the response to Yan or Pnt overexpression. Whereas overexpression of a constitutively active form of Yan (Yan^ACT^ (Rebay and Rubin, 1995)) reduced MHE^WT^ expression 5 fold, only a 3 fold reduction was achieved when the high affinity sites were mutated (Supplemental Fig. S2A). Thus although full repression requires this high affinity ETS pair, at high enough concentrations Yan can interact with weaker sites to repress expression. Overexpression of Pnt increased MHE^WT^ expression 7 fold but only 2-3 fold when ETS sites 2 and/or 3 were mutated (S2B). This suggests that at high concentrations, Pnt will use all ETS sites, and that at normal Pnt concentrations, even when Yan recruitment is compromised by mutation of the ETS 2,3 pair, Pnt remains limiting. Thus relative Pnt and Yan concentrations play important roles in shaping their interaction with the cis-regulatory architecture of the CRE.

Several of our results, namely the expression increases associated with MHE^mut2^, MHE^mut3^, MHE^mut8^, MHE^mut2,3^ and MHE^mut2,5^ (Fig. 2B, 3A, 3D and 3E), contradicted previous findings (Halfon et al., 2000). To rule out the possibility that the 86Fb landing site used in our assays was producing artifacts, we inserted MHE^WT^ and MHE^mut2,3^ into an alternative landing site (attP2), and measured reporter expression. Although not as strong as the increase measured for MHE^mut2,3^ in the 86Fb insertion site, the trend was consistent, with an almost 3-fold increase in expression in the Eve+ clusters and significant ectopic expression outside (Fig. 3F and Supplemental Fig. S3). We conclude that P-element position-effect differences in the earlier study (Halfon et al., 2000) may have masked expression changes that our system more reliably detects.

### The repressive function of the ETS 2,3 pair is conserved across *Drosophila* species

The MHE mutagenesis described above uncovered a pivotal role for the 2,3 high affinity ETS site pair in coordinating repressive inputs. To evaluate the biological relevance of this regulatory capability, we compared ETS syntax in the MHEs of other *Drosophila* species, taking advantage of prior work that aligned *eve* enhancer sequences (Hare et al., 2008) (Fig. 4A,B and Supplemental Fig. S4). Of the three high affinity sites, sites 2 and 3 were conserved over 40 million years of evolution, from *D. melanogaster* to *D. virilis*. The third high affinity site, site 8, was conserved across the *melanogaster* subgroup, but was lost in both the *obscura* and *virilis* groups. Of the lower affinity sites, only site 6, which our reporter analysis had identified as most critical to activation, was conserved across all three groups. Additional ETS sites were noted in the MHE’s of all species outside the *melanogaster* subgroup, but based on our gel shift analysis (Fig. 2A) none of these are predicted to be high affinity since they lack CC at positions -1 and -2. Together these comparisons suggest strong evolutionary pressure both to maintain the unique high affinity 2,3 pair and to limit the binding affinity of other ETS sequences within the enhancer. The correlation between the three sites whose mutation most dramatically altered expression in the MHE-GFP transgenic reporter assay (2,3 and 6) and the three that are most highly conserved suggests our analysis has identified the central determinants of MHE responsiveness to Yan and Pnt.

**Figure 4.**
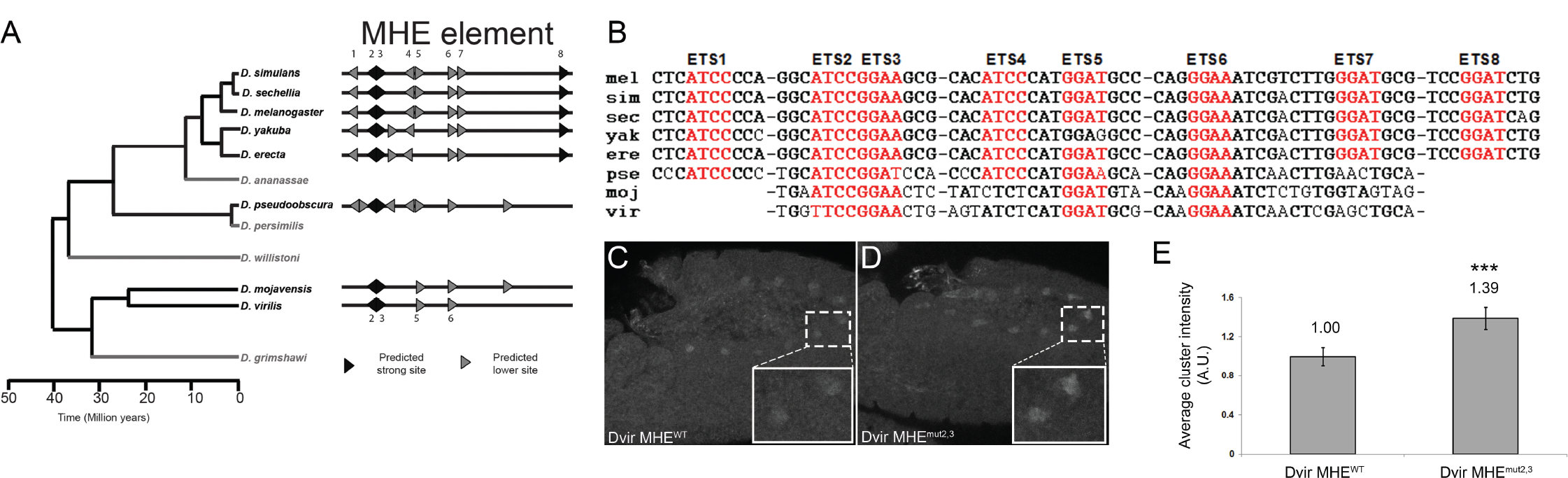
MHE ETS sites 2, 3 and 6 are conserved in distantly related Drosophila species. **A**. Phylogenetic tree (adapted from (Gramates et al., 2017)) and schematic representation of MHE ETS sites in eight different Drosophila species; light gray indicates species whose sequences were not analyzed. Full MHE sequences, as identified by (Hare et al., 2008), are in Supplementary Figure 2. **B**. Comparison of ETS1-8 in the MHE. Base pairs identical to *D. melanogaster* (*mel*) sequences are bolded, the GGAA/T cores are in red, and dashes indicate intervening sequence of various lengths. Additional ETS sites not found in *mel* are not depicted (see Supplemental Figure 2). **C,D**. Stage 11 embryos expressing one copy of *D. virilis* (*Dvir*) MHE^wt^-GFP or *Dvir* MHE^mut2,3^-GFP. Zoomed views of two clusters are shown (white box). **E**. Quantification of the elevated *Dvir* MHE^mut2,3^ reporter expression compared to *Dvir* MHE^WT^, confirms the evolutionary conserved role of the ETS2,3 pair in repression. Error bars show SEM.

To bolster this conclusion, we validated the repressive function of the ETS 2,3 pair in the *D. virilis* MHE using cross-species reporter transgenes. Consistent with previous work (Hare et al., 2008), the control reporter, DvirMHE^WT^, drove GFP expression in the 10 clusters of Eve+ cells (Fig. 4C); expression appeared weaker than that driven by the *melanogaster* MHE^WT^ reporter (Fig. 1B’), presumably reflecting the low degree of overall sequence conservation between these two highly diverged species. Simultaneous mutation of the ETS 2,3 pair increased reporter expression (Fig. 4C-E), emphasizing the functional significance and biological relevance of this high affinity tandem in MHE repression.

### Yan repressive complexes organized by MHE ETS sites 2 and 3 limit access by transcriptional activators

Eve expression in the cardiogenic clusters requires a combination of activating inputs from the mesodermal determinant Twist (Twi), the Ras/MAPK effector Pnt and the Dpp effector Mothers against Dpp (Mad) (Halfon et al., 2000). While the importance of direct competition between Yan and Pnt is well established (Flores et al., 2000; Halfon et al., 2000; Hayashi et al., 2008; Xu et al., 2000), whether Yan repressive complexes also limit access of other transcriptional activators is not known. If so, the increased expression of the MHE^mut2,3^ reporter might reflect not only a shift in the balance of Yan-Pnt competition to favor Pnt, but also increased access to Mad and Twi. To avoid the complications of measuring reporter expression in null mutant backgrounds with cell fate and patterning defects, we tested this hypothesis by asking whether halving the genetic dosage of these three TFs, which does not alter cell fate specification, would moderate the elevated MHE^mut2,3^ reporter expression.

Focusing first on *pnt*, we found that *pnt* heterozygosity reduced MHE^mut2,3^ expression levels by almost 50% (Fig. 5A-C). MHE^mut2,3^ levels were also reduced when either *mad* or *twi* gene dosages were halved (Fig. 5A, D, E). The control experiments showed that expression driven by the MHE^WT^ reporter in *pnt*/+ or *mad*/+ embryos was not statistically different from that measured in a wild type background, while a modest but significant decrease was measured in the *twi*/+ embryos (Fig. 5A). Taken together, these results suggest that Pnt, Mad and Twi all contribute to the increased and ectopic expression measured for the MHE^mut2,3^ reporter. Focusing on Mad and Twi, loss of the Yan-repressive complexes normally organized by the high affinity ETS pair could either relieve a steric block to Mad/Twi occupancy or remove a quenching function that normally dampens the activating output from the assembled Mad/Twi complexes.

**Figure 5.**
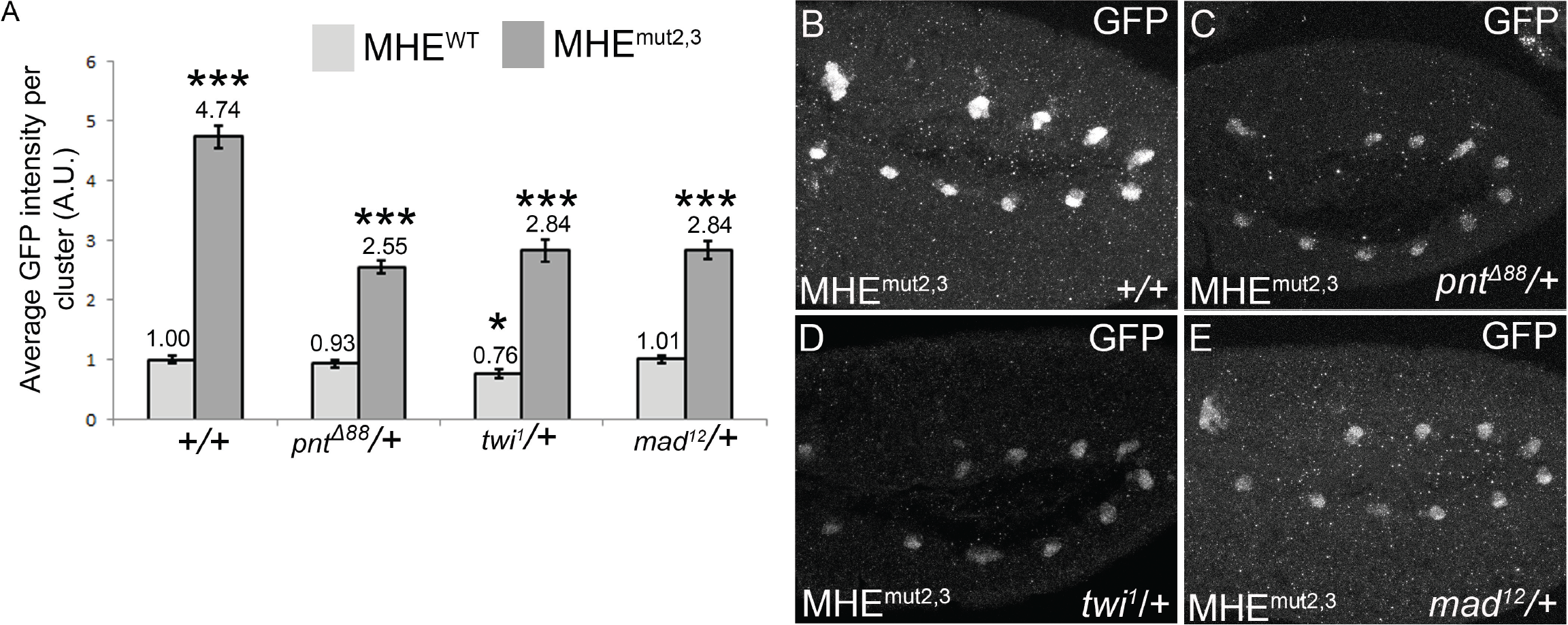
Pointed, Mad and Twist all contribute to the elevated expression of the MHE^mut2,3^ reporter. **A**. Quantification and comparison of average MHE^WT^ (light gray bars) versus MHE^mut2,3^ (dark gray bars) GFP reporter expression intensity in wild type (+/+) versus *pnt, twi* or *mad* heterozygotes. MHE^mut2,3^ reporter expression dropped by about 40-45% in all three heterozygotes, with the greatest response to *pnt*, MHE^WT^ levels were not statistically different in *pnt* or *mad* heterozygotes, but were reduced by about 25% in *twi*/+ embryos. Statistical significance after Bonferroni correction from either MHE^WT^ in the +/+ background for light gray bars and from MHE^mut2,3^ in +/+ for dark grays, with the exception of MHE^mut2,3^ in +/+ which was compared to MHE^WT^ in the same background; *, p <0.05: **, p <0.01; and ***, p <0.001. Error bars show SEM. **B-E**. Representative Stage 11 embryos showing the reduced MHE^mut2,3^ reporter expression in *pnt*/+ (C), *twi*/+(D) or *mad*/+ (**E**) embryos.

### Yan can bind cooperatively to paired ETS sites in the MHE, with affinity and spacing between sites most important for effective repression

Our mutational analysis highlighted the unique importance of ETS sites 2 and 3 to Yan repression at the MHE. Although *in vitro* studies of the cooperative DNA binding ability of the human ortholog of Yan, TEL1, found that ETS site orientation and spacing differences up to 55bp were irrelevant (Green et al., 2010), the significance of such syntax variation to repressive mechanism is not known. To gain insight, we performed additional mutagenesis and re-engineering of the MHE cis-regulatory architecture.

First, we confirmed that Yan can bind cooperatively to paired ETS sites, and that the high affinity 2,3 pair can recruit Yan more effectively than the lower affinity 4,5 and 6,7 pairs. Our substrate was a radiolabeled DNA probe containing the two ETS consensus sites used by Green et al. (Green et al., 2010). Yan monomers bound independently to the two sites, producing at low concentrations, a faster migrating complex corresponding to a single bound monomer and at increasing concentrations, a slower migrating complex corresponding to two bound monomers (Fig. 6A). Incubation of the probe with Yan dimers produced the slower migrating complex, even at low protein concentrations, consistent with cooperative binding (Fig. 6B). Mutation of one site in the probe limited formation of the slower migrating complex, confirming that dimerization favors ternary complexes in which both Yan molecules bind DNA (Fig. 6B). Analogous results were obtained using probes containing pairs of MHE-derived ETS sites, with the slower migrating complex predominant even at low Yan concentrations for the high affinity 2,3 pair but only apparent at higher concentrations for the lower affinity 4,5 and 6,7 pairs (Fig. 6C-E). Thus, Yan can bind cooperatively to DNA sequences derived from the MHE, and the high affinity of sites 2,3 permits full occupancy even at low Yan concentration.

**Figure 6.**
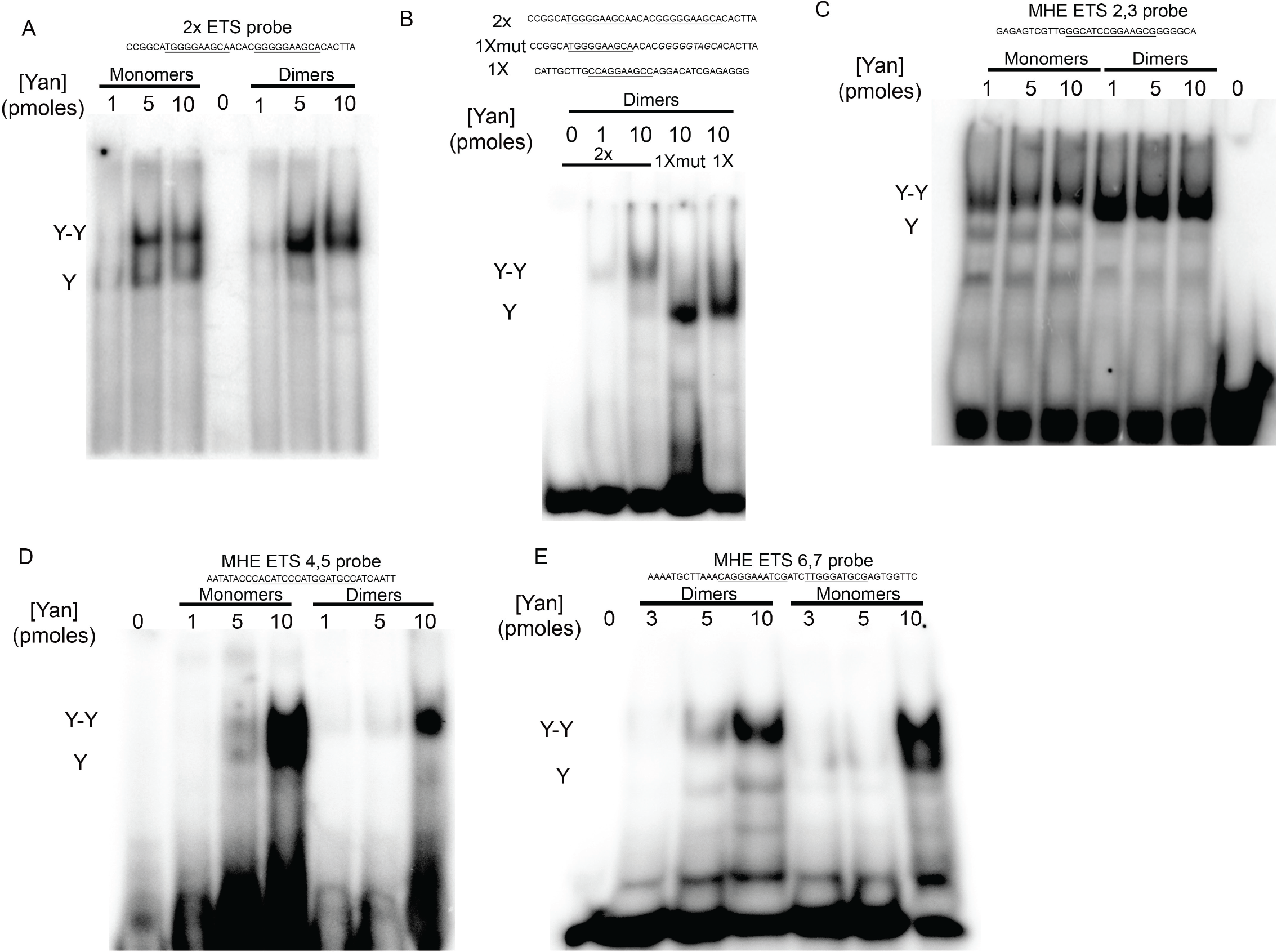
Yan binds cooperatively to tandem ETS pairs derived from the MHE. **A-E**. Gel shifts with Yan monomers (Yan^A86D^) or Yan dimers (1:1 Yan^A86D^:Yan^V105R^). Increasing concentrations of total Yan protein are indicated above each gel. **A**. Cooperative binding by the dimer to a probe with two consensus ETS sites (2x ETS) results in only the slower migrating species at the higher concentrations, whereas both single-bound (Y) and double-bound (Y-Y) species remain evident with Yan monomer. **B**. Mutation of one of the ETS sites in the 2x ETS probe (1xmut), or using a probe with a single site (1x) results in only the faster migrating species (Y) when incubated with Yan dimers, indicating that cooperative DNA binding requires two functional ETS sites. **C-E**. Cooperative binding to probes derived from the MHE. With the high affinity 2,3 pair, the lowest concentrations of Yan dimer fully shift to the Y-Y species whereas higher concentrations are required with the lower affinity 4,5 and 6,7 pairs.

We next exploited features of the endogenous MHE syntax to test the importance of site affinity, orientation and spacing to effective transcriptional repression at paired ETS sites. If high-affinity Yan binding underlies the unique contribution of the 2,3 pair to MHE expression, then introducing a new high-affinity tandem elsewhere in the enhancer should restore repression of the MHE^mut2,3^ reporter. To test this idea, we added a ninth ETS site adjacent to site 8 to form a new pair that closely matched the endogenous 2,3 configuration. We anticipated that this manipulation would be both minimally disruptive to the enhancer and most readily interpretable because although site 8 is subject to Yan-Pnt competition (Fig. 3A, F), repression dominates (Fig 2B). Gel shifts confirmed strong Yan binding affinity and cooperativity (Fig. 7A-B). Quantification of GFP reporter expression showed that the newly engineered 9,8 ETS pair suppressed the elevated expression associated with MHE^mut2,3^ back to MHE^WT^ levels (Fig. 7C). This confirms that a high-affinity tandem ETS pair can be sufficient for effective repression.

**Figure 7.**
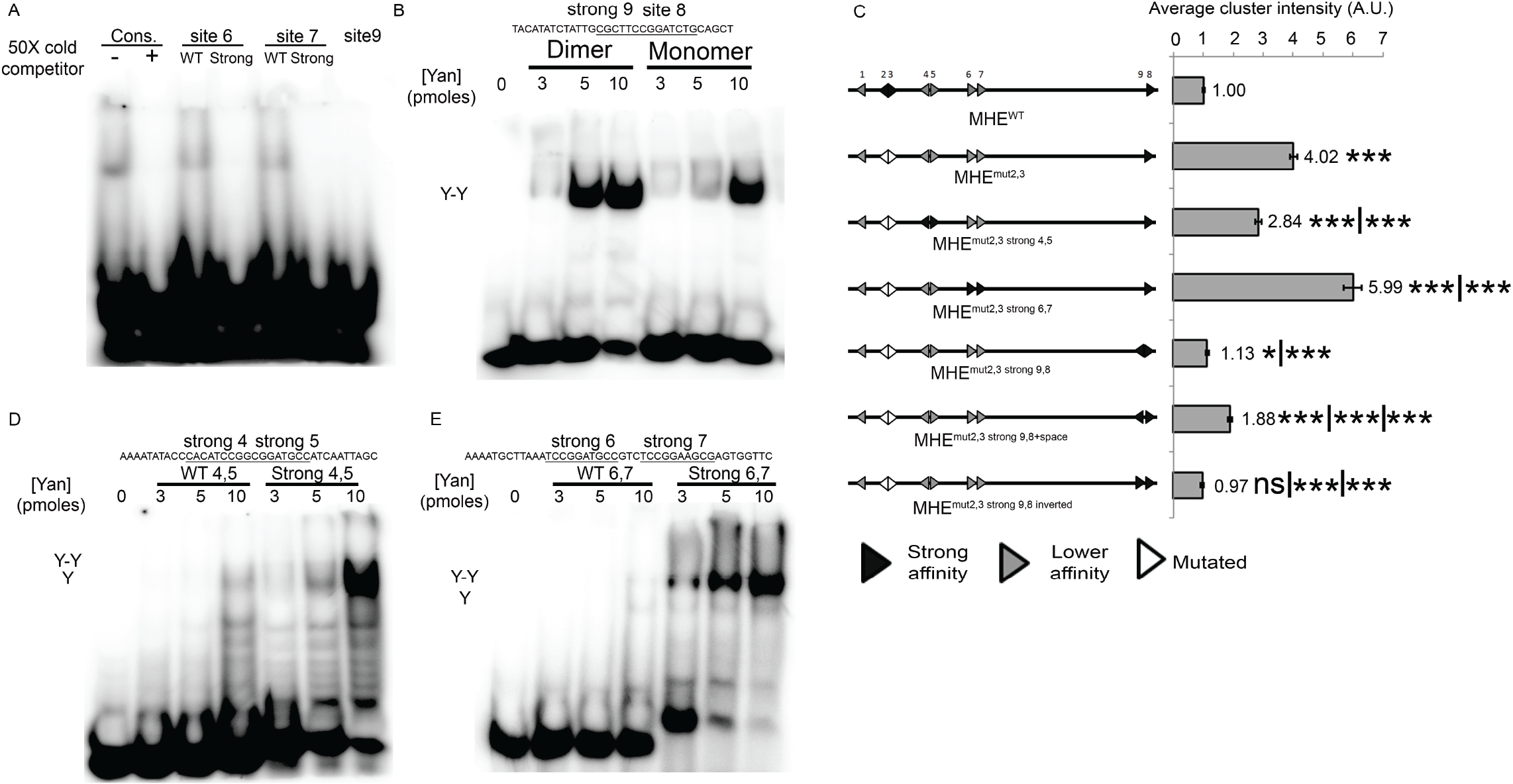
Engineered strong tandem ETS pairs can restore repression of the MHE^mut2,3^ mutant reporter. **A**. Competitive gel shift assay showing that mutation of ETS sites 6 and 7 increased their binding affinity as designed. The added site 9 is also high affinity. **B**. Gel shift assay showing more effective recruitment of Yan dimers than monomers to the strong 9,8 ETS pair. **C**. Quantification of average reporter gene cluster intensity shows that with the exception of the strong 6,7 pair that increased expression, introducing strong ETS pairs into the MHE^mut2,3^ element reduced expression. Error bars show SEM. Statistical significance after Bonferroni correction from either MHE^WT^, MHE^mut2,3^ or MHE^mut2,3 strong9,8^ (where relevant) are depicted. *, p <0.05: **, p <0.01; and ***, p <0.001. **D-E**. Gel shift assays showing increased affinity of the Yan dimers for the engineered “strong” ETS4,5 and 6,7 pairs relative to the wild type (WT) counterparts.

Manipulation of the two other endogenous ETS pairs in the MHE allowed us to investigate the importance of ETS site orientation and spacing. Focusing first on orientation, we mutated the ETS sites in the 6,7 pair toward the high affinity sequences of sites 2 and 3 but maintained their parallel orientation; EMSAs confirmed the desired increase in Yan binding strength and cooperativity (Fig. 7A, 7E). However not only was the 6,7 high affinity pair unable to restore repression, levels of MHE^mut2,3+strong6,7^ were actually higher than those of MHE^mut2,3^, raising the possibility that a parallel ETS site orientation is incompatible with effective repression. Arguing against this, when we inverted sites 9 and 8 to put them in parallel, repression was equivalent to that achieved with the initial anti-parallel configuration (Fig 7C). We conclude that effective Yan repression can be organized by high affinity ETS tandems regardless of site orientation, and that some other property of the 6,7 pair confers its unique contribution to activation of MHE expression. One possibility is that disruption of an annotated binding site for Ladybird, a known repressor of *eve*, that falls between sites 6 and 7 (Liu et al., 2008)(Supplemental Fig. 1a) could be responsible for the increased expression of the MHE^mut2,3+strong6,7^ reporter.

To assess whether spacing between paired ETS sites impacts repression we turned to the 4,5 pair where three bases separate the two GGAT cores; this is in contrast to the 2,3 pair or its engineered 9,8 mimic where the GGAA/T cores are immediately juxtaposed. Gel shifts demonstrated that the mutagenesis designed to increase the affinity of sites 4,5 (see Methods) increased cooperative Yan binding (Fig. 7D) and consistent with higher affinity sites favoring Yan repression, expression of the MHE^mut2,3+strong4,5^ reporter was reduced relative to MHE^mut2,3^ (Fig 7C). However, reporter expression was not repressed back to the MHE^WT^ baseline as it had been with the introduced 9,8 ETS pair, suggesting the spacing might be reducing effectiveness of Yan repression. Because the MHE sequence did not permit removal of this 3bp spacer without reducing binding strength of the 4,5 pair, to test this conclusion further we instead added a 4bp spacer to the engineered 9,8 tandem. The resulting two-fold increase in reporter expression is consistent with the increased spacing reducing the effectiveness of Yan repression (Fig 7C). Keeping in mind the formal caveat that one can never rule out unexpected responses when manipulating enhancer sequence, these results suggest that in contrast to *in vitro* DNA binding behavior where cooperative binding appears agnostic to site orientation or spacing (Green et al., 2010), *in vivo*, increased spacing between paired high affinity ETS sites can reduce the effectiveness of repression.

### MHE ETS sites 2 and 3 organize Yan repressive complexes across the *eve* locus to confer robustness to expression

Having established the importance of the ETS site 2,3 pair to MHE reporter expression, we next used CRISPR/Cas9 to explore its contribution in the context of the whole locus. To control for the impact of genetic background, we first re-engineered the wild type MHE to create an *eve*^*MHEWT*^ allele (see Methods and Supplemental Fig. S5-S7). Specification of Eve+ cells and viability were normal, but average Eve levels per cluster in *eve*^*MHEWT*^ homozygotes were reduced relative to *w*^*1118*^ embryos (Fig. 8A, 8E and Supplemental Fig. S5A). Although the sequence changes used to introduce the PAM sites are outside the minimal enhancer and do not impact known TF binding sites (Supplemental Fig. S7), they could alter unknown activating sites or interactions. Alternatively, the reduction in Eve levels could reflect more complicated influences of genetic background (Jiang et al., 2015). With this in mind, we used *eve*^*MHEWT*^ as the reference background for all subsequent analyses.

**Figure 8.**
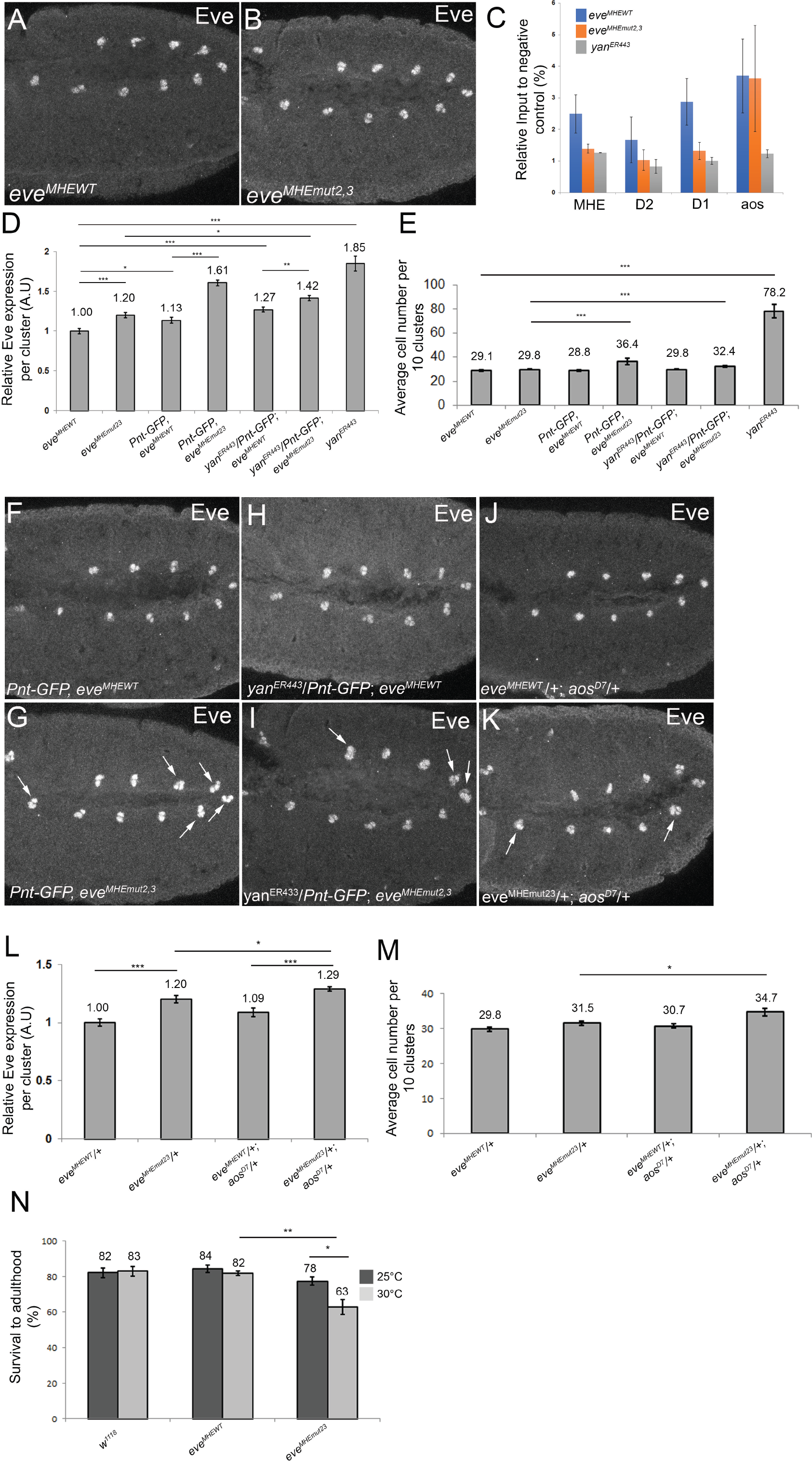
*eve^mut2,3^* embryos have elevated Eve expression, reduced Yan occupancy, and increased sensitivity to genetic and environmental stress, highlighting the importance of ETS2,3-mediated regulation to the robustness of cardiac cell fate specification. **A, B**. Representative Stage 11 embryos expressing two copies of *eve^MHEWT^* or *eve^MHEmut2,3^* and stained for Eve. **C**. ChIP-qPCR from stage 11 embryos was performed at the MHE and two other CREs at the *eve* locus, *D1* and *D2*, and one CRE from *aos*. Signals were normalized to a negative CRE control (see Methods). Reduced occupancy was measured at all three *eve* CREs, but not at *aos*, in *eve^MHEmut2,3HDR^*. Error bars show SEM. **D**. Quantification of average Eve levels per cluster in different genetic backgrounds. Error bars show SEM. Statistical significance after Bonferroni correction is depicted with *, p <0.05: **, p <0.01; and ***, p <0.001. **E**. Average number of Eve+ cells per 10 clusters in the same genetic backgrounds as in (**D**) show that specification of ectopic Eve+ cells occurs in the *eve^MHEmut2,3^* background when the Pnt:Yan ratio is increased. Error bars show SEM. **F-K**. Representative Stage 11 embryos stained for Eve in different genetic backgrounds. Embryos in (**F,H**) express two copies of *eve^MHEWT^*, in (**G, I**) two copies of *eve^MHEmut2,3^* and in (**J,K**) a single copy of *eve^MHEWT^* or *eve^MHEmut2,3^*, respectively. Examples of clusters with 4 or more Eve+ cells are indicated with yellow arrowheads in **G, I** and **K. L**. Quantification of average Eve levels per cluster shows that *aos* heterozygosity similarly increases levels in *eve^MHEWT^* and *eve^MHEmut2,3^* embryos. Error bars show SEM. **M**. *aos* heterozygosity increases the average number of Eve+ cells specified in *eve^MHEmut2,3^* embryos. Error bars show SEM. **N**. The reduced survival of *eve^MHEmut2,3^* embryos is enhanced by temperature stress.

We next mutated ETS sites 2 and 3 to generate *eve*^*MHEmut2,3*^. Average Eve expression levels in the mesodermal clusters were 20% higher than in *eve^MHEWT^* control embryos (Fig. 8B, D). Although this is the predicted result of reduced Yan recruitment and repression, the number of Eve+ cells was unchanged and the homozygotes eclosed with normal Mendelian expectations from balanced stocks (Supplemental Table S3). However in survival assays in which stage 11 embryos were counted and placed in vials and then cultured to adulthood, *eve*^*MHEmut2,3*^ appeared less fit than either *eve*^*MHEWT*^ or w^1118^ controls (Fig. 8N). Culturing the embryos at elevated temperature further reduced *eve*^*MHEmut2,3*^ survival but had no effect on the *eve*^*MHEWT*^ or w^1118^ controls (Fig. 8N).

To determine whether the elevated Eve expression and reduced fitness of *eve*^*MHEmut2,3*^ embryos might result from altered Yan-mediated regulation, we analyzed Yan chromatin occupancy by ChIP-qPCR. Yan binding was reduced at the MHE but unchanged at a positive control CRE from the *argos* (*aos*) locus (Fig. 8C). Thus Yan recruitment to the endogenous *eve* MHE is strongly dependent on the ETS 2,3 tandem, exactly as predicted by our *in vitro* and isolated enhancer-reporter analyses.

Our previous work showed that 3D interactions between the MHE and two other CREs, termed D1 and D2, stabilize Yan occupancy and repression at *eve* such that deletion of any one CRE reduced recruitment to the other (Webber et al., 2013a). We therefore asked whether the reduced Yan recruitment to the MHE also impacted Yan binding at the D1 and D2 elements. Remarkably, significant reductions were measured at both D1 and D2, suggesting that the molecular mechanism by which Yan repressive complexes are organized in 3D across the *eve* locus requires Yan’s cooperative recruitment to MHE ETS sites 2 and 3.

The similarity in phenotypic consequence between mutating the ETS2,3 pair and deleting the D1 or D2 CREs, namely reduced Yan occupancy, increased Eve levels but no cell fate defects, and sensitivity to environmental stress, motivated us to explore further parallels. Specifically, our earlier work showed that the addition of genetic stress to a regulatory system already compromised by deletion of the D1 or D2 CRE impacted the specification of Eve+ cells, and ultimately cardiac function and viability (Webber et al., 2013a). We therefore asked whether genetic manipulations that increase the Pnt:Yan ratio, but do not produce phenotypes in wild type embryos, would induce extra Eve+ cells in the compromised repressive background of *eve*^*MHEmut2,3*^.

Three different genetic interactions were tested: increased *pnt* dose, decreased *yan* dose and decreased *argos (aos)* dose. First, using a functional *pnt-GFP* BAC transgene (Boisclair Lachance et al., 2014) to increase Pnt levels, we found that doubling *pnt* dose increased Eve levels and perturbed cell fate specification (Fig. 8D, E, G and Supplemental Fig. S5B). Although removing one copy of *yan* had no significant effect on Eve intensity or Eve+ cell numbers, when we simultaneously reduced *yan* and added one copy of *pnt-GFP*, which on its own had no effect (Fig. 8I), Eve levels increased (Fig. 8D) and extra Eve+ cells were specified (Fig. 8E, 8H). Finally, we halved the dose of *argos* (*aos*), which encodes an EGFR antagonist whose complete loss lead to an expansion in the number of Eve cells similar to what is observed in *yan* mutants or when Pnt is ectopically expressed (Carmena et al., 2002; Halfon et al., 2000). Consistent with the results of directly manipulating Yan and Pnt dose, Eve levels were increased (Fig. 8L) and extra Eve+ cells were specified (Fig. 8M, 8K). Examining the survival of *eve*^*MHEmut2,3*^ embryos to adulthood in the experiments described above revealed a consistent inverse correlation between the extent of ectopic Eve+ cell fate specification and viability (Supplemental Table S3 and Supplemental Fig. 5C), implying that the specification defects observed at this stage ultimately impair cardiac function. In all these experiments, only minor expression increases and no cell fate changes or reduced viability were detected in the *eve*^*MHEWT*^ controls (Fig. 8).

We conclude that the conserved ETS 2,3 pair provides a critical site for installing Yan repression at *eve* to prevent both elevated expression in the newly specified Eve+ signaling cells and ectopic cell fate induction in the immediately surrounding cells. When these sites are mutated, the residual recruitment of Yan to the MHE and other regulatory elements is sufficient to organize the transcriptional dynamics that support normal cell fate specification, but only under ideal conditions. Finally, these results confirm the power of the transgenic reporter analyses to pinpoint relevant features of endogenous *eve* regulation.

## Discussion

Focusing on ETS binding motifs within a conserved regulatory module, the eve muscle heart enhancer (MHE), we have identified a simple syntax that allows for robust qualitative and quantitative control of enhancer output. Based on extensive mutagenesis, we conclude that Pnt-mediated activation of *eve* expression relies on multiple low-affinity ETS sites with little regard for spacing or orientation. In contrast, Yan’s preference for paired strong affinity sites with optimal spacing enables the assembly of repressive complexes that dampen *eve* expression in newly specified cardiac precursors where Yan levels are low and prevent ectopic eve induction in the surrounding mesoderm where levels of activating TFs such as Twist are high. CRISPR/Cas9 mediated mutation of a high affinity ETS pair in the endogenous MHE confirmed the importance of optimized syntax for Yan-mediated repression and uncovered an unexpected role in longer-range organization of Yan complexes across the locus. We speculate that distinct interpretations of ETS syntax may enable more complex interactions between Yan and Pnt at target enhancers than current models assume, and that differential usage of enhancer syntax by repressors and activators could be a widespread strategy to ensure specific yet robust patterns of gene expression. Further, the demonstration that the cis-regulatory logic at an individual enhancer not only influences short range interactions at that enhancer, but can also foster longer-range communication across multiple CREs, uncovers a novel regulatory mechanism that may prove to be broadly relevant.

Our finding that the lower affinity ETS sites at the MHE are primarily dedicated to activation by Pnt adds to the growing awareness of the importance of low affinity transcription factor binding sites for converting thresholds of signaling activity into spatio-temporally appropriate patterns of gene expression (reviewed in (Crocker et al., 2016)). Pnt’s activating inputs at the MHE have long been known to play a key role in determining the dynamics of *eve* induction in response to RTK signaling (Halfon et al., 2000). At stage 11, Pnt is broadly expressed throughout the mesoderm (Boisclair Lachance et al., 2014), yet only three Eve+ cells are specified per cluster. We suggest that preferential use of low-affinity ETS sites ensures Pnt-dependent activation of the MHE only occurs in cells that have been switched to a “high Pnt state” by RTK signaling. Effective MHE activation by Pnt requires multiple sites, as individual mutation of any of the low-affinity sites both reduced reporter gene expression and limited maximal activation by overexpressed Pnt. This suggests that the combinatorial usage of multiple low affinity sites confers significant sensitivity across a wide range of Pnt concentrations. Across Drosophila species, although multiple low affinity ETS sites are predicted, only site 6 is conserved, raising the possibility that Pnt may be fairly agnostic to syntax. Alternatively, because mutation of site 6 produced the greatest drop in MHE expression, despite its slightly higher affinity for Yan in the gel shift assay, perhaps site 6 provides a critical pattern on-switch, with the other low affinity sites serving as buffers that tune the final level of expression. This would explain why the *D. virilis* MHE, predicted to have only two low affinity ETS sites, drives reporter expression in the expected pattern, yet at a lower level than the *D. melanogaster* MHE. Additional manipulation of the MHE, both in reporters from different species and in the whole locus, will be needed to understand how the affinity and position of site 6 within the overall cis-regulatory logic make it so uniquely responsive to Pnt.

In addition to converting signaling thresholds into the correct spatio-temporal domains of expression, binding site affinity can independently influence how transcription factors with similar DNA binding specificity discriminate between sites. For example, in the case of Hox family transcriptional activators, low-affinity sites provide specificity while higher affinity sites permit greater promiscuity (Berger et al., 2008; Crocker et al., 2015; Noyes et al., 2008). Our study uncovers a different use of site affinity in organizing the competing inputs of an ETS family repressor-activator pair, with Yan relying on strong affinity ETS sites, and in particular on a tandem pair of such sites, and Pnt favoring lower affinity. Another repressor-activator pair with analogous high versus low affinity preferences is Cubitus interruptus (Ci), a TF that is converted from a repressor to an activator during Hedgehog signaling. Re-engineering of Ci binding sites from low to high affinity restricts target gene expression to regions of highest Hedgehog signaling, consistent with the revised syntax favoring repression over activation. Using computational modeling, the *in vivo* enhancer expression patterns could only be predicted correctly when cooperative binding was introduced to Ci repression (Parker et al., 2011), an interesting observation in light of our conclusion that cooperative Yan recruitment to tandem ETS sites is critical to repression.

Our previous work exploring the *in vivo* functionality of a Yan protein in which the SAM-SAM interface has been mutated to prevent self-association further supports the importance of repressor cooperativity. Specifically we found that although Yan monomers are recruited to enhancers genome-wide in a pattern close to that of wild type Yan, adequate repression does not occur and phenotypes consistent with *yan* loss of function ensue (Webber et al., 2013a). This work also noted the prevalence of clustered high affinity ETS sites across a number of Yan ChIP targets, suggesting that the mechanisms uncovered in our dissection of MHE ETS site syntax might be broadly applicable. Focusing on *eve*, we suspect that at the resolution of individual ETS sites, in the absence of SAM-mediated cooperativity, Yan occupancy of the ETS2,3 tandem would be insufficiently stable either to compete appropriately with Pnt, and perhaps with Mad and Twi, at the MHE or to organize the necessary 3D communication across the locus. Given the numerous examples of TFs that can serve as both activator and repressor, it will be interesting to explore how broadly low versus high affinity binding site preferences and repressor cooperativity are used to organize the competing regulatory inputs that drive signal-induced cell fate transitions.

We also note a parallel between the consequences of mutating the high affinity ETS2,3 pair in the endogenous *eve* locus and the findings of an earlier analysis in which we deleted three different Yan bound CREs within a genomic Eve-YFP BAC transgene (Webber et al., 2013b). In this earlier study, while deleting the pattern-driving MHE almost completely ablated mesodermal Eve-YFP induction, deleting a “repressive” Yan-bound element (referred to as D1) increased Eve-YFP expression about 1.5 fold and led to the specification of extra Eve+ cells. Additionally, deletion of either the MHE or the D1 in the BAC transgene reduced Yan occupancy, not just at the deleted element, but also at the remaining intact CREs. Here, we report a comparable loss of Yan occupancy across the *eve* locus upon mutation of the MHE ETS2,3 pair, but only a 1.2 fold increase in Eve levels and no cell fate specification defects. The discrepancy between reduced Yan occupancy and increased Eve levels in the *eve*^*MHEmut2,3*^ mutant relative to the D1 deletion mutant suggests that deleting an entire CRE not only compromises Yan occupancy across the locus, but also disrupts additional repressive inputs. Consistent with this interpretation, the *eve*^*MHEmut2,3*^ background appeared highly sensitized, as a two-fold increase in *pnt* dose was sufficient to boost Eve levels to 1.6 fold above wild type and increased the specification of Eve+ cells, almost exactly matching the effects of deleting an entire “repressive” CRE. Further exploration of the how high affinity ETS tandems organize Yan repression at and between CREs, and how this coordinates the competing and collaborating inputs from other TFs will be needed to test these ideas at *eve* and more broadly at other target genes.

To conclude, we propose a working model in which Yan and Pnt’s differential interpretation of ETS syntax adds a “dimmer” capability to the classic on/off switch, thereby refining its sensitivity and tunability. Using *eve* as the example, prior to the onset of RTK-induced cardiac cell fate specification or in cells subject to sub-maximal signaling, we suggest that Yan’s bias for paired high affinity sites ensures an effectively 100% probability of occupancy at those sites, and hence stable repression. In this regime, Yan either also outcompetes Pnt at the lower affinity sites to occupy fully the CRE, or, if Yan levels are limiting, then its preference for high affinity sites and relative “distaste” for lower affinity sites could offer Pnt an opportunity to occasionally occupy the latter and perhaps attenuate Yan repression. In contrast, if Yan and Pnt had identical ETS binding preferences, we would expect a less tuned response to RTK signaling; indeed, when we removed the high affinity 2,3 pair, and hence the strong bias toward Yan occupancy and repression at the MHE, stochastic ectopic expression was induced in the surrounding mesoderm where RTK levels are sub-maximal. Thus their distinct preferences ensure that only maximal RTK activation will trigger the necessary shift in Yan-Pnt occupancy to activate *eve* expression. Further, while previous models assumed a complete switch from total Yan occupancy to total Pnt occupancy as Eve+ cell fates are specified, our work suggests that the high affinity tandem still recruits Yan repressive input, even in Eve+ cells with very low Yan concentration. We speculate that the ability to apply continued Yan repressive input after cell fate induction may contribute to the robustness of certain developmental transitions by stabilizing the newly acquired cell fate. In agreement with this, in the context of the endogenous *eve* locus, disruption of the ETS 2,3 pair sensitized *eve* to both fluctuations in upstream signaling and environmental stress.

More broadly, we speculate that the interplay between the cis-regulatory logic of a CRE and the unique biophysical parameters of different TFs permits evolution to fine-tune gene expression output to a specific threshold depending on each cell’s developmental requirement. In the case of Yan-Pnt regulated genes, the interplay between the degree of Yan self-association and ETS syntax enables this repressor-activator pair to discriminate between ETS sites with unexpected precision. Further, instead of RTK activation inducing a complete switch from Yan occupancy to Pnt occupancy as cell fates are induced, the cooperative recruitment of Yan to paired high affinity sites may enable newly differentiating cells with lower Yan:Pnt ratios to sustain the Yan repressive influence needed to ensure precision and robustness of the gene expression patterns. We suggest these ideas provide an interesting new vantage point for considering how single nucleotide polymorphisms in TF binding sites may heighten susceptibility to disease by compromising robustness of gene regulatory networks.

## Materials and methods

### Drosophila genetics

*Strains used:* The following chromosomes were from the Bloomington *Drosophila* Stock Center: *w^1118^*, *pnt^Δ88^*, *mad^12^*, *twi^1^*, *aos^Δ7^*, *Df(3L)BSC562(aos)*, *UAS-PntP1*, *nos-phiC31;AttP2*, *nos-phiC31;86Fb*, *TM3,Sb-Ser,twist-GAL4,UAS-GFP(TTG)*, *Cyo-twist-GAL4,UASGFP(CTG)*, *TM6B, tubulin-GAL80* and *Cyo,Tub>PBac*.

Additional strains: *yan^E443^* and *yan^E833^* (Karim et al., 1996), *UAS-Yan^ACT^* (Rebay and Rubin, 1995), *Pnt-GFP* (Boisclair Lachance et al., 2014) *vasa-Cas9* (a gift from R. Fehon, GFP and RFP negative), and *eve^MHEWT^* and *eve^MHEmut2,3^* (this study, described below). A list of MHE-pJR20 transgenes (this study) is in Supplementary Table 4.

*CRISPR/Cas9-mediated generation of eve^MHEWT^ and eve^MHEmut2,3^ alleles:* A 2.1 kb region including the wild-type or mutated MHE sequences and mutated PAM sites was inserted in pHD-Scarless (generated by O’Connor-Giles lab, DGRC, #1364). Both templates were confirmed by sequencing. Guide RNAs were subcloned into the pU6–Bbs1 chiRNA plasmid (Gratz et al., 2013) (Addgene 45946). Each template (300ng/ul) and the two guide RNAs (75ng/ul) were injected into a GFP/ RFP negative vasa-Cas9 strain (a gift from Rick Fehon). G_o_ adults were crossed individually to *w^1118^* and transformants were identified by 3X-Pax-RFP expression in the eyes of the F1 progeny. The 3XPax-RFP piggyBac cassette was excised and progeny without RFP expression were crossed to CTG to establish the *eve^MHEWT^* and *eve^MHEmut2,3^* stocks; the desired alleles were confirmed by restriction digest and sequencing. See Supplementary Methods for additional details.

### MHE Reporter Subcloning and Transgenesis

Quik-change mutagenesis (Stratagene) of MHE^WT^-pBluescript (Halfon et al., 2000) was used to modify ETS binding site sequences (Primers listed in Supplementary Table 4). Elements were shuttled into the pJR20 GFP reporter plasmid; (Rister et al., 2015)) as BamH1 fragments and confirmed by sequencing prior to injection. Transgenes were inserted at the 86Fb (Bischof et al., 2007) or attP2 (Groth et al., 2004) landing sites and outcrossed to *w^1118^* to remove the X-linked ϕ31 integrase.

A 484bp D. *virilis* MHE^WT^ was amplified from genomic DNA using primers: 5’-TACTCCGGCGCTCCTCGAGGTTAATGCACCCAGCAGCC-3’ and 5’-GTACCCCGCGGCCGCTAGCGTTTGCAGTGTAGCTGAAATATATGG-3’ and inserted into BamH1 digested pJR20 using Gibson Assembly. A 484bp gene block (Integrated DNA Technologies, see Supplementary Table 4 for sequence) containing D. *virilis* MHE^mut2,3^ was similarly inserted into pJR20.

### Embryo staining and imaging

Embryos were stained and fixed as previously described (Webber et al., 2013b)(see Supplemental Methods for details). Primary antibodies: rabbit anti-GFP (ThermoFisher Scientific, A6455, 1/2000), chicken anti-GFP (Abcam, ab13970, 1/2000), mouse anti-Eve 3C10 (DHSB, 1/10; (Patel et al., 1992), and mouse anti-Yan 8B12 (DHSB, 1/750, (Rebay and Rubin, 1995)). Secondary antibodies (Jackson Immunoresearch, 1/2000): goat anti-mouse Cy3, goat anti-rabbit 488, and donkey anti-Chicken 488. All primary antibodies have been validated for use in Drosophila embryos and are standard for the field.

For GFP reporter expression measurements, the microscope gain was set for embryos carrying one copy of MHE^WT^-GFP in an otherwise wildtype background, and then the same settings were used to image all other samples fixed and stained in parallel. For experiments with the CRISPR alleles, the gain was set up on the *eve^WT^* allele, unless otherwise stated. Images were obtained using either a Zeiss LSM510 or Zeiss LSM880 confocal microscope using 0.45μm optical sections. GFP reporter expression and Eve expression were measured using ImageJ as previously described (Webber et al., 2013b) and normalized to MHE^WT^/+ or *eve^MHEWT^* expression unless otherwise stated. At least 2 independent experiments were analyzed per genotype. Eve expression was quantified similarly to reporter gene expression.

### Statistical analysis

All results were analyzed using 2-way ANOVA analysis to test for statistical significance. When found to be significantly different, a two-way Student T-Test with Bonferroni correction was applied to validate statistical significance in a pairwise manner. For all experiments, at least 40 clusters (4 embryos), but in general 100 clusters (10 embryos) were analyzed.

### In vitro gel shift assays

Recombinant Yan^A86D^ and Yan^V105R^ (aa 1-499) were prepared by TEV protease cleavage of GST-fusion proteins purified from BL21 codon plus *E. coli* cells (See Supplemental Methods for details). Yan protein in the eluate was assessed for quality and concentration by SDS-PAGE followed by ImageJ quantification of the intensity of Yan products. Eluates were stored at 4°C for up to 5 days.

Double-stranded probes were made by End-labeled with T4 polynucleotide kinase (New England Biolabs) and gamma^32^P-ATP (Perkin Elmer) for 1hr at 37°C, prior to annealing. All oligonucleotide sequences are listed in Supplementary Methods.

For competition assays, about 30pmol of Yan^A86D^ was incubated with 50X cold probe for 20 minutes on ice in 25 μl reactions (0.5 μg polydI-dC, 5 mg BSA and 25% glycerol in 13 mM HEPES pH 7.9, 40 mM KCl, 0.7 mM EDTA, 0.3 mM DTT and 1pmol of labelled probe). 100x cold competitor was added and reactions were incubated an additional 20 minutes on ice. Cooperative binding was assessed by incubating equimolar amounts of Yan^V105R^ and Yan^A86D^ on ice for 20 minutes in the presence of 1pmol of labelled probe. Samples were resolved on 10% polyacrylamide gels (competition assay) or on 4-20% gradient gels (cooperative binding) to capture both bound and free probe run in 0.5X Tris-Borate-EDTA (TBE) buffer for 45 minutes at 120 volts.

### ChIP-qPCR

ChIP from stage 11 embryos (5h20-7h20 at 25°C) was performed as described in Webber et al., Genetics, 2013 (see Supplemental Methods for details).

## Acknowledgments

We thank Mark Halfon for pBluescript-MHE^WT^, Jens Rister and Claude Desplan for pJR20, Cynthia Horth, Hitoshi Matakatsu and Rick Fehon for the generation of GFP/RFP-negative *vasa-Cas9* strain and Aaron Mitchell-Dick with help injecting constructs. We acknowledge the Bloomington Drosophila Stock Center (NIH P40OD018537), the Developmental Studies Hybridoma Bank (created by the NICHD of the NIH), and the Drosophila Genomics Resource Center (NIH 2P40OD010949) for critical reagents. We thank Rebay and Fehon lab members for helpful discussions and Trevor Davis, Matt Hope and Richard Mann for comments on the manuscript. This work was supported by NIH R01 GM080372 to I.R. and by the Genomics Core Facility through a University of Chicago Cancer Center Support Grant P30 CA014599.

## References

Berger, M.F., Badis, G., Gehrke, A.R., Talukder, S., Philippakis, A.A., Peña-Castillo, L., Alleyne, T.M., Mnaimneh, S., Botvinnik, O.B., Chan, E.T., et al. (2008). Variation in Homeodomain DNA Binding Revealed by High-Resolution Analysis of Sequence Preferences. Cell 133, 1266–1276.

Bischof, J., Maeda, R.K., Hediger, M., Karch, F., and Basler, K. (2007). An optimized transgenesis system for Drosophila using germ-line-specific phiC31 integrases. Proc. Natl. Acad. Sci. U. S. A. 104, 3312–3317.

Boisclair Lachance, J.F., Peláez, N., Cassidy, J.J., Webber, J.L., Rebay, I., and Carthew, R.W. (2014). A comparative study of pointed and yan expression reveals new complexity to the transcriptional networks downstream of receptor tyrosine kinase signaling. Dev. Biol. 385, 263–278.

Carmena, A., Gisselbrecht, S., Harrison, J., Jiménez, F., and Michelson, A.M. (1998). Combinatorial signaling codes for the progressive determination of cell fates in the Drosophila embryonic mesoderm. Genes Dev. 12, 3910–3922.

Carmena, A., Buff, E., Halfon, M.S., Gisselbrecht, S., Jiménez, F., Baylies, M.K., and Michelson, A.M. (2002). Reciprocal regulatory interactions between the Notch and Ras signaling pathways in the Drosophila embryonic mesoderm. Dev. Biol. 244, 226–242.

Crocker, J., Abe, N., Rinaldi, L., McGregor, A.P., Frankel, N., Wang, S., Alsawadi, A., Valenti, P., Plaza, S., Payre, F., et al. (2015). Low affinity binding site clusters confer HOX specificity and regulatory robustness. Cell 160, 191–203.

Crocker, J., Preger-Ben Noon, E., and Stern, D.L. (2016). The Soft Touch: Low-Affinity Transcription Factor Binding Sites in Development and Evolution. In Current Topics in Developmental Biology, pp. 445–469.

Farley, E.K., Olson, K.M., Zhang, W., Brandt, A.J., Rokhsar, D.S., and Levine, M.S. (2015). Suboptimization of developmental enhancers. Science 350, 325–328.

Farley, E.K., Olson, K.M., Zhang, W., Rokhsar, D.S., and Levine, M.S. (2016). Syntax compensates for poor binding sites to encode tissue specificity of developmental enhancers. Proc. Natl. Acad. Sci. U. S. A. 113, 6508–6513.

Flores, G. V, Duan, H., Yan, H., Nagaraj, R., Fu, W., Zou, Y., Noll, M., and Banerjee, U. (2000). Combinatorial signaling in the specification of unique cell fates. Cell 103, 75–85.

Gramates, L.S., Marygold, S.J., Santos, G. dos, Urbano, J.-M., Antonazzo, G., Matthews, B.B., Rey, A.J., Tabone, C.J., Crosby, M.A., Emmert, D.B., et al. (2017). FlyBase at 25: looking to the future. Nucleic Acids Res. 45, D663–D671.

Gratz, S.J., Cummings, A.M., Nguyen, J.N., Hamm, D.C., Donohue, L.K., Harrison, M.M., Wildonger, J., and O’connor-Giles, K.M. (2013). Genome engineering of Drosophila with the CRISPR RNA-guided Cas9 nuclease. Genetics 194, 1029–1035.

Green, S.M., Coyne, H.J., McIntosh, L.P., and Graves, B.J. (2010). DNA binding by the ETS protein TEL (ETV6) is regulated by autoinhibition and self-association. J. Biol. Chem. 285, 18496–18504.

Groth, A.C., Fish, M., Nusse, R., and Calos, M.P. (2004). Construction of Transgenic Drosophila by Using the Site-Specific Integrase from Phage ??C31. Genetics 166, 1775–1782.

Halfon, M.S., Carmena, A., Gisselbrecht, S., Sackerson, C.M., Jiménez, F., Baylies, M.K., and Michelson, A.M. (2000). Ras pathway specificity is determined by the integration of multiple signal-activated and tissue-restricted transcription factors. Cell 103, 63–74.

Hare, E.E., Peterson, B.K., Iyer, V.N., Meier, R., and Eisen, M.B. (2008). Sepsid even-skipped enhancers are functionally conserved in Drosophila despite lack of sequence conservation. PLoS Genet. 4, e1000106.

Hayashi, T., Xu, C., and Carthew, R.W. (2008). Cell-type-specific transcription of prospero is controlled by combinatorial signaling in the Drosophila eye. Development 135, 2787–2796.

Hollenhorst, P.C., McIntosh, L.P., and Graves, B.J. (2011). Genomic and biochemical insights into the specificity of ETS transcription factors. Annu. Rev. Biochem. 80, 437–471.

Hope, C.M., Rebay, I., and Reinitz, J. (2017). DNA Occupancy of Polymerizing Transcription Factors: A Chemical Model of the ETS Family Factor Yan. Biophys. J. 112, 180–192.

Inukai, S., Kock, K.H., and Bulyk, M.L. (2017). Transcription factor-DNA binding: beyond binding site motifs. Curr. Opin. Genet. Dev. 43, 110–119.

Jiang, P., Ludwig, M.Z., Kreitman, M., and Reinitz, J. (2015). Natural variation of the expression pattern of the segmentation gene even-skipped in melanogaster. Dev. Biol. 405, 173–181.

Karim, F.D., Chang, H.C., Therrien, M., Wassarman, D.A., Laverty, T., and Rubin, G.M. (1996). A screen for genes that function downstream of Ras1 during Drosophila eye development. Genetics 143, 315–329.

Klämbt, C. (1993). The Drosophila gene pointed encodes two ETS-like proteins which are involved in the development of the midline glial cells. Development 117, 163–176.

Knirr, S., and Frasch, M. (2001). Molecular Integration of Inductive and Mesoderm-Intrinsic Inputs Governs even-skipped Enhancer Activity in a Subset of Pericardial and Dorsal Muscle Progenitors. Dev. Biol. 238, 13–26.

Liu, J., Qian, L., Han, Z., Wu, X., and Bodmer, R. (2008). Spatial specificity of mesodermal even-skipped expression relies on multiple repressor sites. Dev. Biol. 313, 876–886.

Mackereth, C.D., Schärpf, M., Gentile, L.N., MacIntosh, S.E., Slupsky, C.M., and McIntosh, L.P. (2004). Diversity in structure and function of the Ets family PNT domains. J. Mol. Biol. 342, 1249–1264.

Nitta, K.R., Jolma, A., Yin, Y., Morgunova, E., Kivioja, T., Akhtar, J., Hens, K., Toivonen, J., Deplancke, B., Furlong, E.E.M., et al. (2015). Conservation of transcription factor binding specificities across 600 million years of bilateria evolution. Elife 4.

Noyes, M.B., Christensen, R.G., Wakabayashi, A., Stormo, G.D., Brodsky, M.H., and Wolfe, S.A. (2008). Analysis of Homeodomain Specificities Allows the Family-wide Prediction of Preferred Recognition Sites. Cell 133, 1277–1289.

O’Neill, E.M., Rebay, I., Tjian, R., and Rubin, G.M. (1994). The activities of two Ets-related transcription factors required for drosophila eye development are modulated by the Ras/MAPK pathway. Cell 78, 137–147.

Oldridge, D.A., Wood, A.C., Weichert-Leahey, N., Crimmins, I., Sussman, R., Winter, C., McDaniel, L.D., Diamond, M., Hart, L.S., Zhu, S., et al. (2015). Genetic predisposition to neuroblastoma mediated by a LMO1 super-enhancer polymorphism. Nature 528, 418–421.

Parker, D.S., White, M.A., Ramos, A.I., Cohen, B.A., Barolo, S., Barolo, S., Posakony, J.W., Ashe, H.L., Briscoe, J., Wang, Q.T., et al. (2011). The cis-regulatory logic of Hedgehog gradient responses: key roles for gli binding affinity, competition, and cooperativity. Sci. Signal. 4, ra38.

Patel, N.H., Ball, E.E., and Goodman, C.S. (1992). Changing role of even-skipped during the evolution of insect pattern formation. Nature 357, 339–342.

Qiao, F., Song, H., Kim, C.A., Sawaya, M.R., Hunter, J.B., Gingery, M., Rebay, I., Courey, A.J., and Bowie, J.U. (2004). Derepression by Depolymerization. Cell 118, 163–173.

Rebay, I., and Rubin, G.M. (1995). Yan functions as a general inhibitor of differentiation and is negatively regulated by activation of the Ras1/MAPK pathway. Cell 81, 857–866.

Rister, J., Razzaq, A., Boodram, P., Desai, N., Tsanis, C., Chen, H., Jukam, D., and Desplan, C. (2015). Single-base pair differences in a shared motif determine differential Rhodopsin expression. Science (80-.). 350, 1258–1261.

Scholz, H., Deatrick, J., Klaes, A., and Klämbt, C. (1993). Genetic dissection of pointed, a Drosophila gene encoding two ETS-related proteins. Genetics 135, 455–468.

Siggers, T., and Gordân, R. (2014). Protein-DNA binding: Complexities and multi-protein codes. Nucleic Acids Res. 42, 2099–2111.

Slupsky, C.M., Gentile, L.N., Donaldson, L.W., Mackereth, C.D., Seidel, J.J., Graves, B.J., and McIntosh, L.P. (1998). Structure of the Ets-1 pointed domain and mitogen-activated protein kinase phosphorylation site. Proc. Natl. Acad. Sci. U. S. A. 95, 12129–12134.

Soldner, F., Stelzer, Y., Shivalila, C.S., Abraham, B.J., Latourelle, J.C., Barrasa, M.I., Goldmann, J., Myers, R.H., Young, R.A., and Jaenisch, R. (2016). Parkinson-associated risk variant in distal enhancer of α-synuclein modulates target gene expression. Nature 533, 1–20.

Sopko, R., and Perrimon, N. (2013). Receptor Tyrosine Kinases in Drosophila Development. Cold Spring Harb. Perspect. Biol. 5, a009050–a009050.

Webber, J.L., Zhang, J., Cote, L., Vivekanand, P., Ni, X., Zhou, J., Nègre, N., Carthew, R.W., White, K.P., and Rebay, I. (2013a). The relationship between long-range chromatin occupancy and polymerization of the drosophila ets family transcriptional repressor yan. Genetics 193, 633–649.

Webber, J.L., Zhang, J., Mitchell-Dick, A., and Rebay, I. (2013b). 3D chromatin interactions organize Yan chromatin occupancy and repression at the even-skipped locus. Genes Dev. 27, 2293–2298.

Wei, G.-H., Badis, G., Berger, M.F., Kivioja, T., Palin, K., Enge, M., Bonke, M., Jolma, A., Varjosalo, M., Gehrke, A.R., et al. (2010). Genome-wide analysis of ETS-family DNA-binding in vitro and in vivo. EMBO J. 29, 2147–2160.

Xu, C., Kauffmann, R.C., Zhang, J., Kladny, S., and Carthew, R.W. (2000). Overlapping activators and repressors delimit transcriptional response to receptor tyrosine kinase signals in the Drosophila eye. Cell 103, 87–97.

Zhang, J., Graham, T.G.W., Vivekanand, P., Cote, L., Cetera, M., and Rebay, I. (2010). Sterile alpha motif domain-mediated self-association plays an essential role in modulating the activity of the Drosophila ETS family transcriptional repressor Yan. Mol. Cell. Biol. 30, 1158–1170.

Zhu, L.J., Christensen, R.G., Kazemian, M., Hull, C.J., Enuameh, M.S., Basciotta, M.D., Brasefield, J.A., Zhu, C., Asriyan, Y., Lapointe, D.S., et al. (2011). FlyFactorSurvey: a database of Drosophila transcription factor binding specificities determined using the bacterial one-hybrid system. Nucleic Acids Res. 39, D111–D117.

